# Molecular evolution and inheritance pattern of Sox gene family among Bovidae

**DOI:** 10.1101/2022.08.19.504581

**Authors:** Mabel O. Akinyemi, Muyiwa S. Adegbaju, Anastasia Grytsay, Osamede H Osaiyuwu, Jessica Finucan, Ibukun M. Ogunade, Sunday O. Peters, Bolaji N. Thomas, Olanrewaju B. Morenikeji

## Abstract

Sox gene is an evolutionarily conserved family of transcription factors that play important roles in cellular differentiation and numerous complex developmental processes. In vertebrates, Sox proteins are required for cell fate decision, morphogenesis, and control self-renewal in embryonic and adult stem cells. Sox gene family has been well studied in multiple species including humans but scanty or no study in Bovidae. In this study, we conducted a detailed evolutionary analysis of this gene family in Bovidae, including their physicochemical properties, biological functions, and patterns of inheritance. We performed a genome-wide cataloguing to explore the Sox gene family using multiple bioinformatics tools. Our analysis revealed conserved motifs that are crucial to the ability of Sox genes to interact with the regulatory regions of target genes and orchestrate multiple developmental and physiological processes. Importantly, we report a unique motif being EFDQYL/ELDQYL found in SoxE and SoxF groups. Further analysis revealed that this motif sequence accounts for the binding and transactivation potential of Sox proteins. Protein-protein interaction showed significant interaction among Sox genes and related genes implicated in embryonic development and the regulation of cell differentiation. We conclude that Sox gene family uniquely evolved among Bovidae with a few exhibiting important motifs that drives several developmental and physiological processes.

## Introduction

The Sox genes belong to a large group of genes in which the DNA binding domain is called a high mobility group (HMG), encoding diverse and well-conserved transcription factors, and consisting of 20 genes in vertebrates and only a handful in invertebrates (Lefebvre, 2019, Stevanovic et al., 2021). Sox genes share homology with the HMG box of sex-determining region Y (Sry) gene, a founder member of the Sox gene family, required to specify male phenotype. This group of genes owns their acronym to sharing of 46% or more identity to the Sry gene in the HMG box (Dy et al., 2008).

Vertebrate Sox genes have been divided into eight groups (A-H) based on conserved protein domains and similarity in their amino acid sequences (Schepers et al., 2002, Stevanovic et al., 2021). Members of the same group are highly similar to each other within and outside the HMG box, while those of different groups share a lower degree of identity in the HMG box and no significant identity outside the domain (Dy et al., 2008). Most members of this group are scattered throughout the genome with the exception of Sry and Sox3, which are located on autosomes (Stevanovic et al., 2021).

Sox protein control transcription in several ways, the protein domain contacts the minor groove of DNA to AACAAAG, AACAAT, and related sequences and induces a sharp bend of DNA thereby allowing the protein to contribute an important structural role in the assembly of transcription enhancer complexes (Dy et al., 2008). They exhibit their structural role by shaping the regulatory regions and establishing physical contacts between transcription factors bound to the same target gene promoter or enhancer (Stevanovic, 2021). Thus, the regulatory functions of Sox proteins require the cooperation of interacting partners that binds DNA and enable specific selection of target genes (Kendoh et al., 2010). This type of cooperation is dynamic, allowing Sox proteins to regulate different events by changing partner factors: a key factor that drives the progression of developmental processes (Kondoh and Kamachi 2010).

All Sox genes have a specific expression pattern and most play important roles in the determination of cell fate and differentiation of cells into specific lineages, such as embryonic stem cells, neuronal and glial cells, Sertoli cells, chondrocytes, etc (Dy et al., 2008). They have also been shown to be capable of reprogramming differentiated somatic cells into pluripotent stem cells (Takahashi et al., 2006, Takahashi et al., 2007), activating genes that are important for maintaining the pluripotent cell state (Takahashi and Yamanaka 2016), and induction of delta crystallin gene in the lens (Smith et al., 2009).

Functions of the members of the Sox B1(Sox) family overlap although each has a function during the migration of neuronal precursors in the ganglionic eminence (Ekonomou et al., 2005). Sox2 and Sox3 deficiency cause pituitary defects (Rizzoti et al., 2004) Single knockouts of Sox 5 or Sox6 in mice lead to some skeletal abnormalities, and a Sox5 and Sox6 double knockout leads to a lack of chondrogenesis (Smits et al., 2001). Mutations in Sox18 induce severe cardiovascular and hair follicle defects and lymphatic dysfunction (Pennisi et al., 2000; Kamachi and Kondoh, 2013). Mutational disruptions in Sox9 lead to the failure of otic placode invagination (Barrionuevo et al., 2008). Members of the SoxF subfamily exhibit overlapping functions (Sakamoto et al., 2007) and are expressed in the endothelial cells and their mutation has been reported in lymphatic defects (Francois et al., 2008). Sox18 has been reported to be involved in lymphatic development, its mutation in humans leads to lymphatic dysfunction (Kamachi and Kondoh, 2013). Kanai-Azuma et al 2002 also reported defects in gut tube formation in Sox17 knockout mice. Members of the SoxC subfamily (Sox4, 11, and 12) are expressed in neural and mesenchymal progenitor cells. Mutations in these subfamily lead to malformation of various organs. Sox4 and Sox11 knockout mice exhibit multiple organ defects (Sock et al., 2004). Several studies aimed at discovering more sox protein functions have been impaired by extensive functional redundancy among members of the same group and pleiotropy (Kamachi and Kondoh, 2013) The Bovidae family includes several economically and socially important ruminant species such as cattle, sheep, goats, and antelope, comprising more than 140 recognized species distributed among 49 genera. The evolution of the different members of this family was molded by several mechanisms including temperature adaptations, feeding ecology, vegetation physiognomy, climate fluctuations, immigration, adaptive radiations, and mass extinctions (Matthee and Davis, 2001. Published reports examining the evolutionary history of the Bovidae through morphological data, allozymes, serum immunology, DNA sequence, and mitochondria DNA analyses exists (Escuidero et al 2019). Protein and DNA-based sequence-based methods utilize sequence variations and analysis of conserved domains, in addition to computational methods utilizing genetic variation within and between amino acid sequences to predict functional and structural outcomes. (Hepp et al., 2015; Morenikeji and Thomas, 2019; Singh et al., 2020; Soremekun et al., 2020).

Most of the computational analyses on Sox genes have focused on individual genes and their regulatory functions rather than the gene family. Despite the abundance of information on their functions Sox in various developmental processes, we found no published report on Sox gene family in Bovidae. Taking this into consideration and the fact that Sox genes are significant players in the regulation of developmental processes, we examined the evolutionary dynamics of the Sox gene family within Bovidae by evaluating the biochemical properties, structural prediction, conserved domain analysis, protein-protein interaction, and phylogenetic relationships.

## Materials and Methods

### Sequence retrieval and multiple sequence alignment

Sox genes sequences of the Bovidae family namely: and *Bos taurus* (Cattle), *Bos grunniens* (Domestic Yak), *Bison bison* (Bison), *Capra hircus* (Goat), *Ovis aries* (Sheep), *Bubalus bubalis* (Buffalo), *Bos mutus* (Wild Yak), *Antilocapra americana* (Antelope) species were employed for comparative analysis. All sequences were retrieved in the FASTA format from the National Centre for Biotechnology Information (NCBI) database (https://www.ncbi.nlm.nih.gov/). The Bos sp accession numbers are as follows: SoxA (ABY19364.1), SoxB (XP_005214070.1, NP_001098933.1, XP_027390716.1), SoxB2(NP_001157253.1,XP_024855919.1}, SoxC (NP_001071596.1,XP_024855327.1,XP_010809935.2), SoxD (AAI14659.1,XP_027419183.1, XP_027419995.1) SoxE (XP_002698019.3,XP_027374751.1, XP_027396627.1), SoxF (XP_027404803.1, XP_027416795.1, XP_027415027.1), SoxG (XP_027374313.1), SoxH (XP_027401619). The total number of Sox genes, category and abundance of each subfamily were collated for further analysis.

Sequence alignment was performed using the ClustalW algorithm on the MEGAX software (Kumar et al., 2018). The alignment was imported to itol for phylogenetic tree construction using Maximum Likelihood Tree method, as described (Cicarelli et al., 2006). Thereafter, MEME (Multiple Em for Motif Elicitation) version 5.0.2 (Bailey et al., 2009) was used to predict the conserved motif structures encoded across all Bos Sox genes. All sequences were scanned for the presence of the sequence motif RPMNAFMVW, which has been reported to be conserved for all Sox sequences other than Sry and only sequences that have the motif were included in further analysis. We used Multalin software to identify the conserved amino acid sequence among the species. The sequence logo of the identified domain in the Sox protein family was constructed with WebLogo (http://weblogo.berkeley.edu/logo.cgi)

### Physicochemical characterization of Sox gene family

To decipher the physicochemical properties of Sox gene family in Bovidae, we used Expasy ProtParam server (https://web.expasy.org/protparam/) to compute the molecular weight, isoelectric point (pI), instability index, aliphatic index, and grand average of hydropathicity (GRAVY) of all the proteins in the Bovidae Sox family, as previously described (Morenikeji and Thomas, 2019). The chromosomal location of each Sox gene was retrieved from Unitprot/KB/Swiss-Prot database in NCBI (https://www.ncbi.nlm.nih.gov/)

### Identification of interacting proteins, functional enrichment and pathway analysis

Protein-protein interactions were predicted using the Search Tool for the Retrieval of Interacting Gene database (STRING https://stringdb.org). This is important to elucidate the association of Sox proteins with other molecules. Functional enrichment analysis was performed with PANTHER (http://www.pantherdb.org/) (Mi et al., 2013) and Database for Annotation, Visualization, and Integrated Discovery (DAVID) to classify the genes according to their function, annotated with ontology terms: biological processes, cellular components, and molecular functions of the studied genes.

### Evolutionary phylogenetic Analysis of Sox gene family

To study the evolutionary origin of Sox genes in Bos sp, multiple sequence alignment was performed using ClustalW tool in MEGA11(Kumar et al., 2018) with default parameters, all positions containing gaps and missing data were eleiminated. The phylogenetic trees of Sox gene family in Bos sp were constructed by adopting the Neighbor joining method of MEGA11 software with the following parameters: Poisson correction, pairwise deletion and 1000 bootstrap replicates. The constructed tree files were visualized using *itol* (Interactive Tree of Life, https://itol.embl.de)

## Results

### Genome wide analysis of Sox genes in Bovidae

A total of 350 variants of *SRY*, 49 variants of Sox 5 and 63 variants of Sox 6 were reported in *Bos taurus* (Figure 1). *Bos mutus* has 33 variants of Sox A and 3 variants of Sox 6. The buffalo (*Bubalus bubalis*) has 226 variants of Sox A and 34 variants of Sox 6, while 24 variants of Sox A was reported for Yak. Sheep (*Ovis aries*), has 163 variants of Sox A and 20 variants of Sox 6 while goat (*Capra hircus*) presented with 4 variants of Sox A and 11 variants of Sox 6. *Bison bison* has 77 variants of Sox A and 26 variants of Sox 6. There was no record of Sox variants for antelopes.

**Figure 1:**
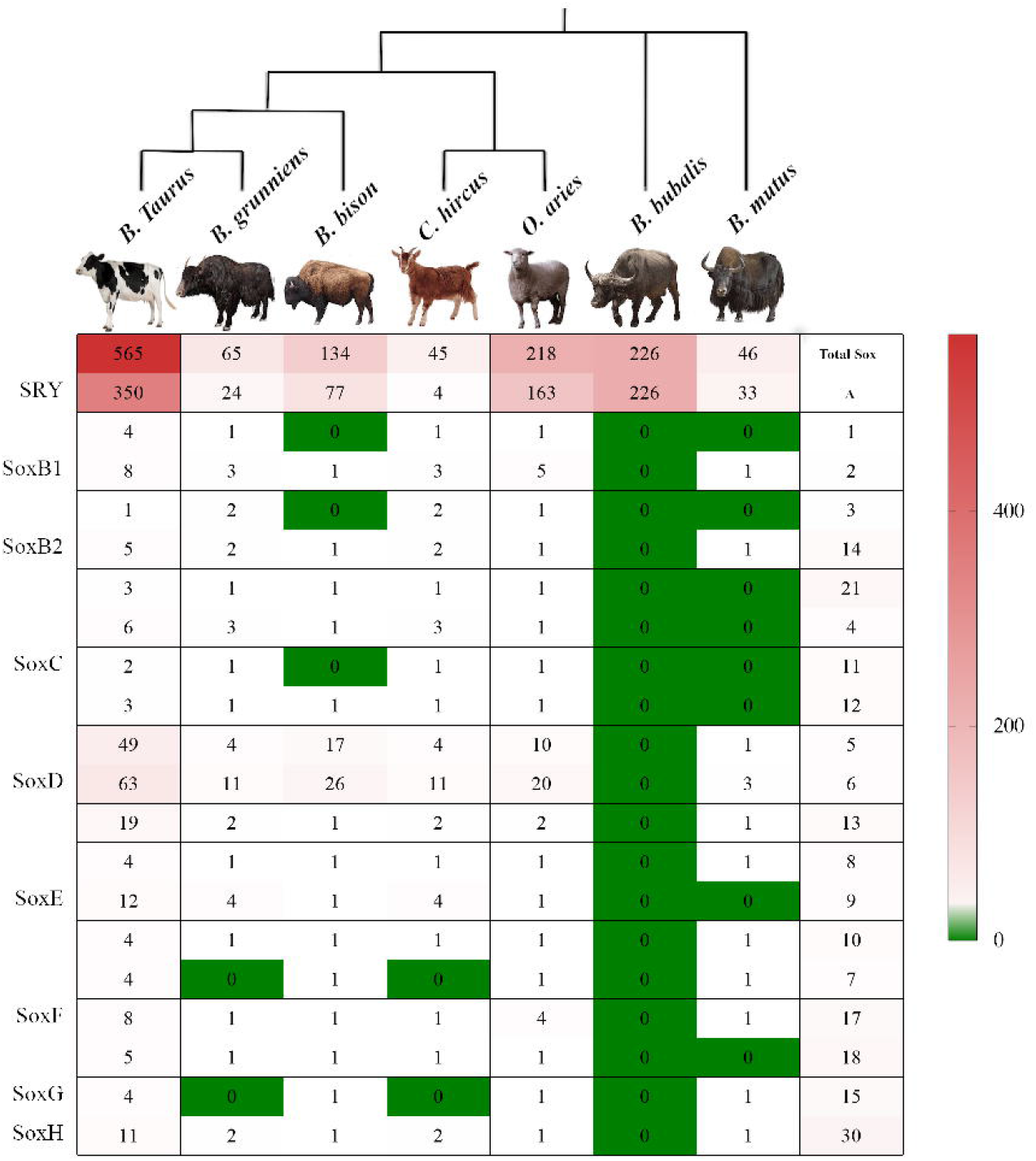
The phylogenetic analysis, categories and abundance of the Sox gene family in Bovidae

### Physicochemical properties of Sox genes

Our analysis of the physicochemical properties reveal that Sox15 had the lowest molecular weight of 25140 KDa. while the Sox D family (5, 6, and 13) had the highest molecular weight values (Table 1). The theoretical isoelectric point(*pI*) is the pH at which a particular molecule carries no net electric charge, this value is useful for understanding the protein charge stability. Overall, the pI ranged from 4.95 to 9.85 (Table 1). Results for instability index reveal that the genes were all unstable (II>40), with the lowest of 41.96 in Sox1 and the highest of 80.78 in Sox9. The aliphatic index, regarded as the factor determining thermostability of globular proteins was lowest in Sox1 (44.44) and highest in Sox13 (71.88). High extinction coefficient and low negative GRAVY values were also observed.

**Table 1.** Physicochemical properties of Bos Sox genes. MW (Molecular weight in KiloDaltons), pI (Isoelectric point), AI (Aliphatic index), II (Instability Index), GRAVY (Grand Average of Hydropathicity Index)

|  | Gene symbol | Accession number | Size | MW (Da) | pI | Chromosome | AI | II | GRAVY | Extinction Coefficient |
| --- | --- | --- | --- | --- | --- | --- | --- | --- | --- | --- |
| soxA | SRY | ABY19364.1 | 229 | 26627.11 | 9.53 | Y | 62.62 | 60.71 | -0.858 | 30285 |
| soxB<sub>1</sub> | Sox1 | XP_005214070.1 | 369 |  |  | 1 | 44.44 | 41.96 | -0.472 | 35870 |
|  | Sox2 | NP_001098933.1 | 320 | 34480.95 | 9.74 | 1 | 48.25 | 59.43 | -0.739 | 37360 |
|  | Sox3 | XP_027390716.1 | 451 | 45113.00 | 9.78 | X | 61.18 | 69.35 | -0.314 | 38850 |
| soxB<sub>2</sub> | Sox14 | NP_001157253.1 | 240 | 26485.39 | 9.68 | 1 | 63.17 | 53.51 | -0.585 | 33015 |
|  | Sox21 | XP_024855919.1 | 277 | 28666.92 | 9.74 | 12 | 69.13 | 58.51 | -0.208 | 31525 |
| soxC | Sox4 | NP_001071596.1 | 481 | 47785.11 | 7.20 | 23 | 54.22 | 58.41 | -0.478 | 35660 |
|  | Sox11 | XP_024855327.1 | 451 | 47147.03 | 4.95 | 11 | 58.16 | 65.27 | -0.652 | 37150 |
|  | Sox12 | XP_010809935.2 | 314 | 33997.76 | 5.14 | 13 | 49.87 | 67.74 | -0.982 | 40575 |
| soxD | Sox5 | AAI14659.1 | 728 | 80127.28 | 6.13 | 5 | 69.48 | 63.71 | -0.746 | 43445 |
|  | Sox6 | XP_027419183.1 | 858 | 95541.56 | 7.93 | 15 | 64.52 | 61.11 | -0.794 | 64790 |
|  | Sox13 | XP_027419995.1 | 623 | 69061.04 | 6.14 | 16 | 71.88 | 69.67 | -0.742 | 41870 |
| soxE | Sox8 | XP_002698019.3 | 534 | 56239.99 | 7.77 | 25 | 52.25 | 59.49 | -0.708 | 63175 |
|  | Sox9 | XP_027374751.1 | 524 | 57173.49 | 6.31 | 19 | 46.98 | 80.78 | -0.984 | 55935 |
|  | Sox10 | XP_027396627.1 | 469 | 50020.20 | 6.19 | 5 | 53.30 | 58.52 | -0.822 | 54445 |
| soxF | Sox7 | XP_027404803.1 | 534 | 57216.30 | 9.62 | 8 | 55.66 | 63.44 | -0.834 | 58830 |
|  | Sox17 | XP_027416795.1 | 410 | 43060.76 | 5.91 | 14 | 60.63 | 65.47 | -0.509 | 39225 |
|  | Sox18 | XP_027415027.1 | 389 | 41337.00 | 8.42 | 13 | 62.96 | 78.07 | -0.573 | 40255 |
| soxG | Sox15 | XP_027374313.1 | 233 | 25140.18 | 9.85 | 19 | 49.06 | 69.49 | -0.875 | 38515 |
| soxH | Sox30 | XP_027401619.1 | 757 | 82746.76 | 8.62 | 7 | 70.01 | 68.76 | -0.603 | 48415 |

Sox 1 Sequence contains several consecutive undefined AA.Its pI and Mw cannot be computed

### Evolutionary Dynamics and phylogenetic analysis

Furthermore, this study examined the evolutionary pattern of 31 Sox sequences of the Bos sp. The multiple sequence alignment indicates that Lysine (K), Proline (P), Arginine R), Glycine (G), and Leucine (L) are conserved in the Sox family (Figure 2). The sequences demonstrated significant variability in both percentage identity and similarity in Bos despite their common evolutionary origin. Analysis of percentage identity and similarity reveals that Sox14 and 21 (71.73%) were highly similar while Sox 7 and 30 exhibited the lowest similarity (15.17%) (Table 2)

**Table 2.**
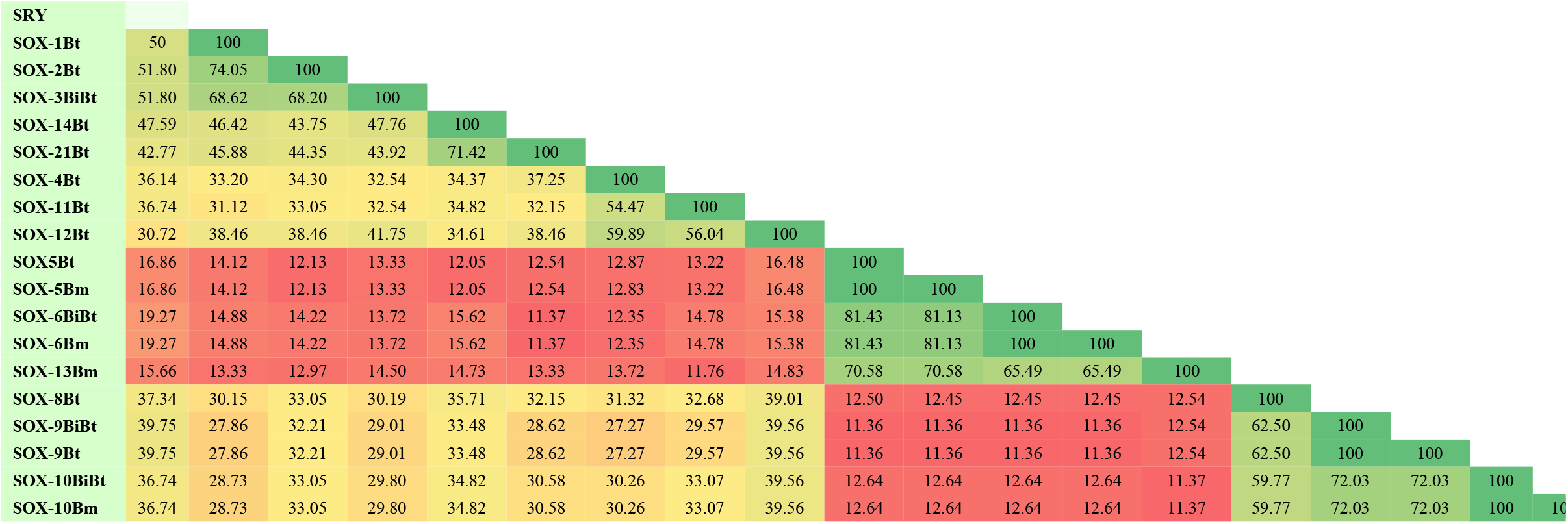
Molecular similarity index of Sox gene family in Bovidae

**Figure 2:**
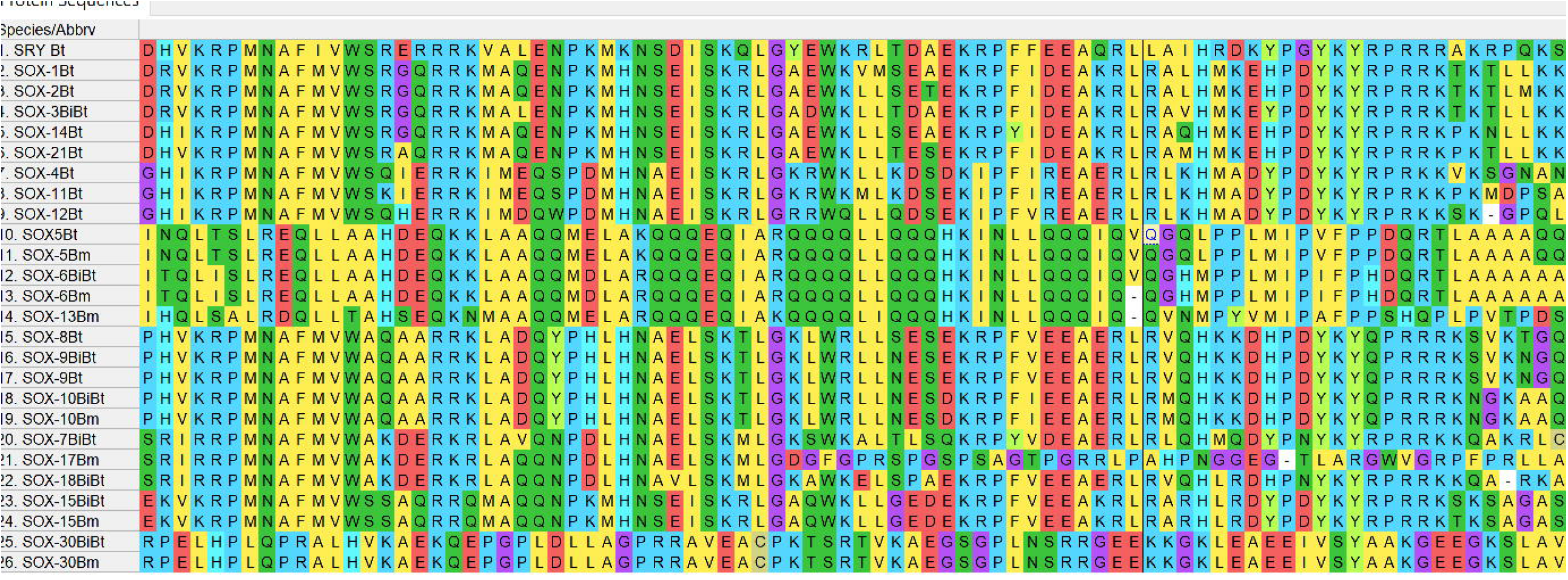
Multiple sequence alignment of Sox genes in Bos sp

Using the distance-based method, we examined the phylogenetic relationships among Bos Sox sequences. A phylogenetic tree based on the alignment showed that the Sox proteins are segregated into nine specific groups: A (*SRY*), B_1_ (Sox1,2 and 3), B_2_ (Sox14 and 21), C (Sox4, 11 and 12), D (Sox5, 6, and 13), E (Sox8, 9 and 10), F (Sox7, 17 and 18), Sox15 and 30 are sole members of Group G and H respectively (Figure 3a). The branch lengths, representative of the extent of divergence suggest a monophyletic arrangement, with Sox B_1_ diverging first. The result is further represented in an unrooted tree, with the expected clustering pattern was observed for all groups (Figure 3b). Figure 4 shows the MSA of the homology of the HMG domain across Sox genes showing that Lysine (K), Proline (P), Glutamate (E), Arginine (R), and Leucine (L) are 100% conserved in this region. The sequence logo showing the relative frequencies of each of the conserved domains and their respective positions is displayed (Figure 5).

**Figure 3:**
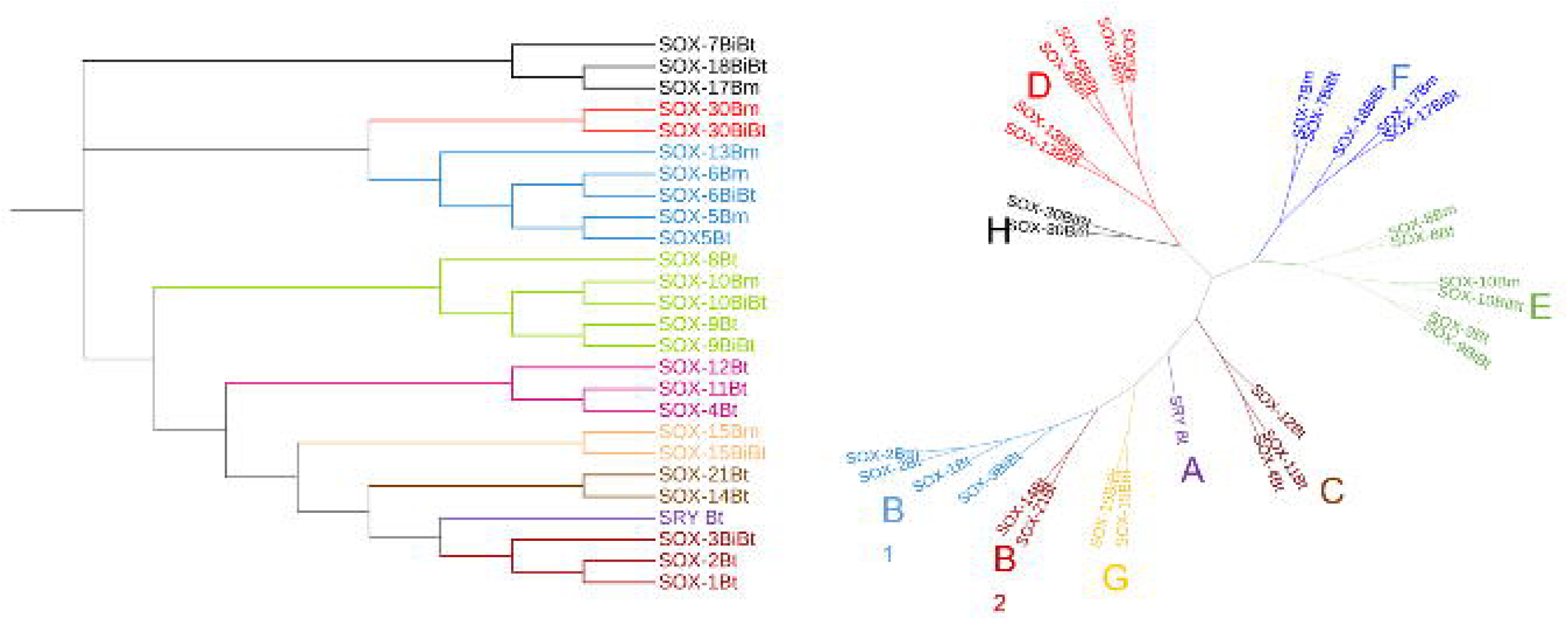
Unrooted phylogeny of Sox genes in Bos sp computed using the neighbor joining method (MEGA). The interactive tree of life (ITOL) online software (https://itol.embl.de) was used to assign color to the different subfamilies

**Figure 4:**
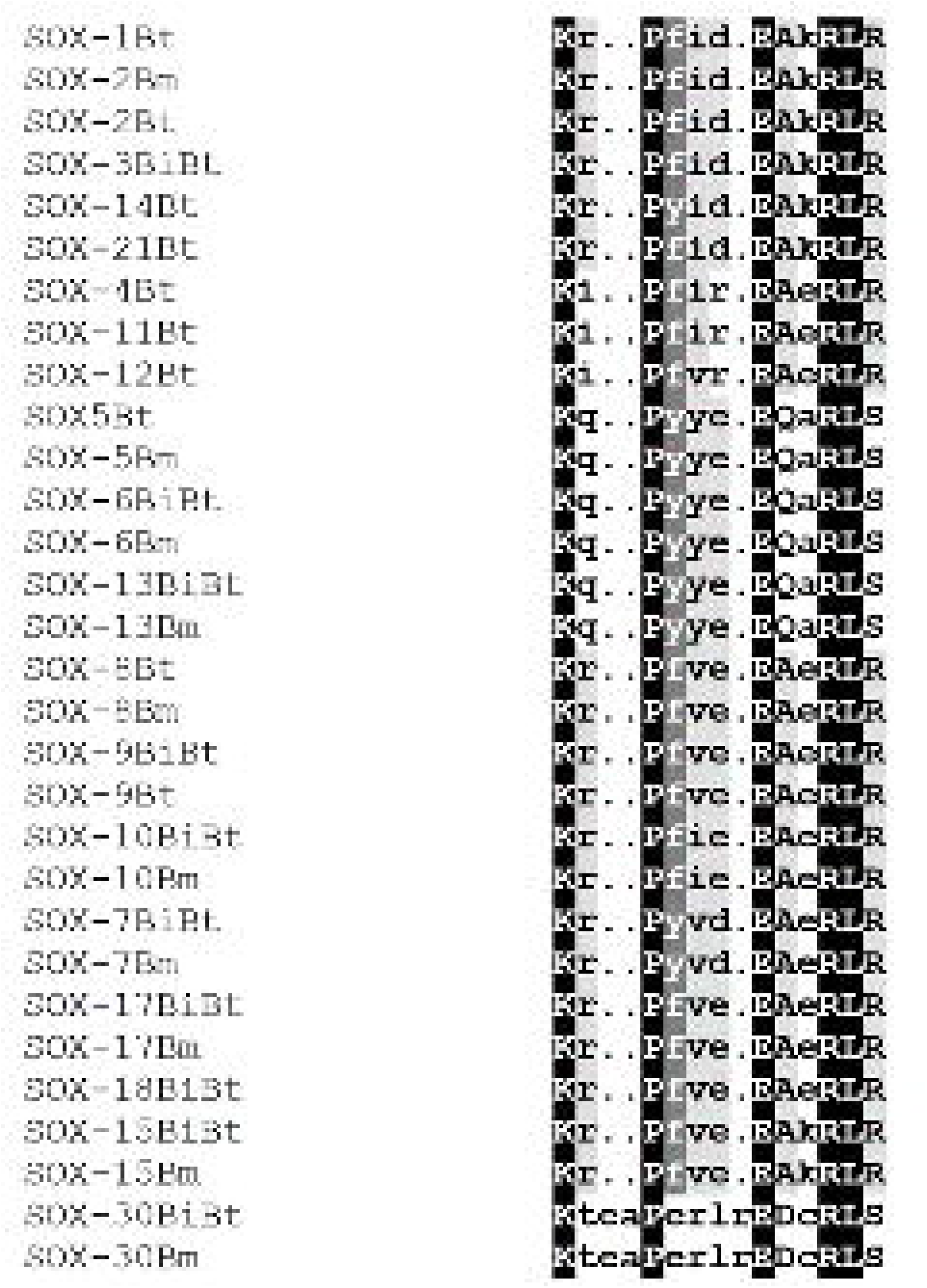
Conserved domain pattern in Sox genes across Bos sp

**Figure 5:**
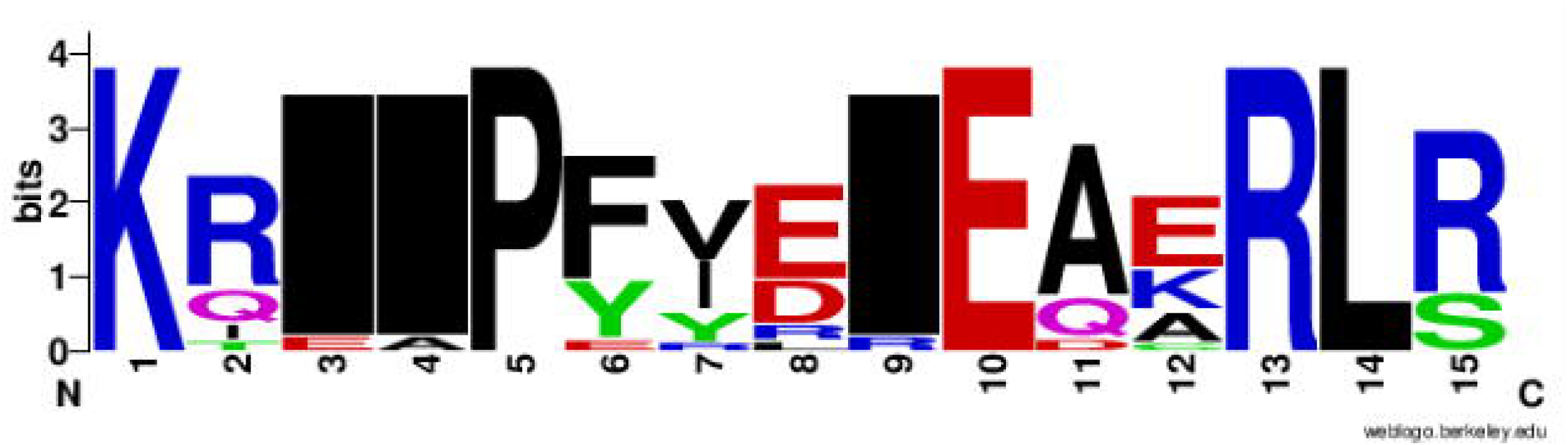
Logo plots of Sox protein sequences displaying the most conserved domain and positions of amino acids. A total of 26 multiple sequence alignments are represented using Weblogo (http://weblogo.berkeley.edu/logo.cgir). the relative frequency of the amino acids is shown on the *y*-axis

### Conserved motif analysis of Sox genes

To further characterize the evolutionary pattern of Sox gene in Bos species, conserved motif analysis was performed using the MEME program. The results indicate that 10 conserved motifs are present in Bos species. Interestingly, we found that conserved motifs 1 and 2 (M1 and M2) exist in all members of the Sox family except Sox 17 which had motif 1 only. The other nine motifs were present in the different Sox groups. M1 and M2 are the core HMG box containing 79 amino acid residues. As shown, five motifs are present in SoxD: Sox13 and SoxE: Sox9, 10, and seven motifs in SoxD: Sox5, (Figure 6a). Four motifs existed in SoxE: Sox8 and three motifs in Sox H: Sox 30. The three-dimensional structure of M1 and M2 of Bos species revealed an L-shaped fold (Figure 6b).

**Figure 6:**
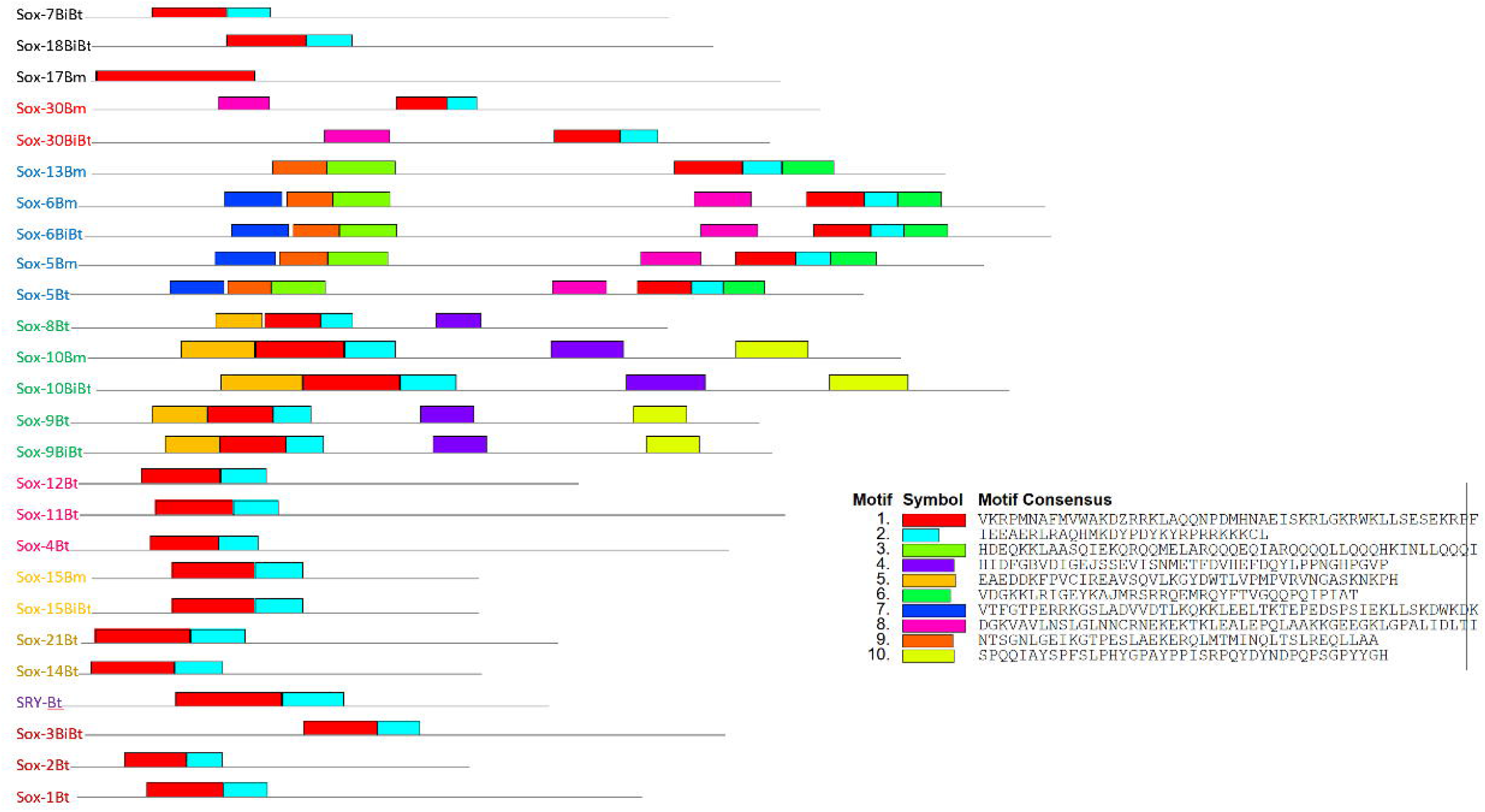
Motif patterns of Bos Sox gene family. Ten differentially conserved motifs are indicated with different colored boxes using the online Multiple Expectation Maximization for Motifs Elicitation (MEME, http://alternate.meme-suite.org) program

### Characterization of functional motifs

All the Sox sequences analyzed were individually scanned for matches against the PROSITE collection of protein signature databases. We found two main domains HMG box B and Sox C terminal domains (Figure 7), with varying frequency across the Sox gene family in Bovidae. We found one proline-rich domain in Sox 9, 17, 18, 15, and 30. The amino acid composition analysis reveals that these proteins have the highest proline concentration among the studied Sox proteins (Sox9-16.98%; Sox17-16.34%; Sox18-19.28%; Sox15-14.59%; Sox30-14.40%) (Table 3).

**Table 3.** Amino acid composition of Sox genes. Ala: Alanine, Cys: Cysteine, Asp: Aspartic Acid, Glu: Glutamic acid, Phe: Phenylalanine, Gly: Glycine, His: Histidine, Ile: Isoleucine, Lys: Lysine, Leu: Leucine, Met: Methionine, Asn: Asparagine, Pro: Proline, Gln: Glutamine, Arg: Arginine, Ser: Serine, Thr: Threonine, Val: Valine, Trp: Tryptophan, Tyr: Tyrosine

|  | Ala | Cys | Asp | Glu | Phe | Gly | His | Ile | Lys | Leu | Met | Asn | Pro | Gln | Arg | Ser | Thr | Val | Trp | Tyr | Total |
| --- | --- | --- | --- | --- | --- | --- | --- | --- | --- | --- | --- | --- | --- | --- | --- | --- | --- | --- | --- | --- | --- |
| SRYBt | 6.14 | 2.63 | 5.26 | 5.7 | 3.51 | 2.63 | 3.95 | 3.07 | 7.02 | 7.89 | 2.19 | 3.95 | 5.26 | 5.7 | 9.21 | 10.96 | 5.26 | 4.82 | 1.32 | 3.51 | 228 |
| SOX-1Bt | 18.97 | 0.27 | 2.17 | 3.52 | 0.54 | 15.45 | 5.96 | 1.36 | 4.34 | 5.96 | 4.61 | 2.98 | 8.4 | 3.79 | 4.61 | 7.59 | 1.63 | 3.25 | 0.81 | 3.79 | 369 |
| SOX-2Bm | 7.67 | 1.47 | 2.95 | 5.6 | 0.29 | 10.32 | 3.54 | 1.47 | 5.31 | 6.78 | 5.9 | 3.54 | 7.67 | 4.13 | 7.67 | 11.5 | 4.13 | 4.13 | 2.06 | 3.83 | 339 |
| SOX-2Bt | 8.13 | 0.31 | 2.81 | 3.75 | 0.63 | 11.88 | 3.44 | 1.25 | 5.31 | 6.25 | 7.81 | 4.69 | 8.13 | 5.63 | 5.31 | 11.25 | 4.38 | 3.75 | 0.94 | 4.38 | 320 |
| SOX-3BiBt | 20.18 | 0.22 | 3.55 | 2.22 | 0.89 | 12.64 | 2.44 | 1.11 | 3.77 | 7.1 | 3.99 | 2.22 | 12.42 | 3.1 | 5.1 | 7.98 | 3.99 | 3.1 | 0.67 | 3.33 | 451 |
| SOX-14Bt | 10.83 | 1.25 | 3.75 | 3.75 | 2.5 | 5.83 | 3.75 | 2.5 | 7.5 | 8.75 | 3.75 | 2.92 | 11.25 | 3.33 | 5.42 | 9.17 | 5 | 2.92 | 1.25 | 4.58 | 240 |
| SOX-21Bt | 22.38 | 0.72 | 2.53 | 3.97 | 2.53 | 8.66 | 3.61 | 1.81 | 6.86 | 8.3 | 3.61 | 2.17 | 10.11 | 1.81 | 4.33 | 7.94 | 1.44 | 2.53 | 1.08 | 3.61 | 277 |
| SOX-4Bt | 14.55 | 0.83 | 4.99 | 4.57 | 2.49 | 13.72 | 2.49 | 2.08 | 6.03 | 6.24 | 1.66 | 2.49 | 6.24 | 1.25 | 3.53 | 19.54 | 2.08 | 2.49 | 0.83 | 1.87 | 481 |
| SOX-11Bt | 11.75 | 1.11 | 7.76 | 7.1 | 2.66 | 11.53 | 1.55 | 2 | 6.87 | 7.76 | 2.44 | 1.55 | 7.76 | 3.55 | 3.99 | 12.64 | 2 | 2.88 | 0.89 | 2.22 | 451 |
| SOX-12Bt | 10.83 | 0.96 | 6.37 | 11.46 | 1.91 | 12.42 | 1.59 | 1.91 | 4.46 | 5.73 | 2.55 | 0.64 | 12.74 | 2.55 | 9.55 | 5.1 | 2.55 | 3.18 | 1.91 | 1.59 | 314 |
| SOX5Bt | 8.52 | 0.41 | 4.53 | 8.24 | 1.65 | 6.87 | 2.2 | 4.12 | 7.01 | 8.24 | 3.57 | 3.57 | 7.83 | 8.38 | 4.67 | 8.79 | 4.12 | 4.4 | 0.41 | 2.47 | 728 |
| SOX-5Bm | 8.16 | 0.53 | 4.14 | 8.02 | 1.74 | 6.95 | 2.54 | 3.88 | 6.95 | 8.02 | 3.48 | 3.74 | 7.75 | 8.42 | 4.68 | 9.22 | 4.55 | 4.41 | 0.4 | 2.41 | 748 |
| SOX-6BiBt | 7.5 | 0.89 | 4.57 | 7.37 | 1.78 | 5.84 | 1.78 | 4.19 | 6.61 | 7.75 | 3.94 | 3.81 | 7.37 | 8.89 | 5.21 | 10.04 | 6.1 | 3.43 | 0.64 | 2.29 | 787 |
| SOX-6Bm | 7.02 | 0.83 | 4.52 | 7.37 | 1.78 | 6.54 | 2.5 | 4.16 | 6.66 | 7.85 | 3.8 | 3.8 | 7.13 | 8.68 | 5.47 | 9.87 | 5.83 | 3.33 | 0.59 | 2.26 | 841 |
| SOX-13BiBt | 7.48 | 1.46 | 4.72 | 8.13 | 1.3 | 5.2 | 2.76 | 2.76 | 6.02 | 9.76 | 2.93 | 3.09 | 9.92 | 8.46 | 5.53 | 9.43 | 3.25 | 5.2 | 0.49 | 2.11 | 615 |
| SOX-13Bm | 7.36 | 1.6 | 5.12 | 7.84 | 1.28 | 5.28 | 2.72 | 2.88 | 5.92 | 9.6 | 2.88 | 3.04 | 9.76 | 8.64 | 5.6 | 9.44 | 3.2 | 5.28 | 0.48 | 2.08 | 625 |
| SOX-8Bt | 12.17 | 1.12 | 5.24 | 4.31 | 1.69 | 10.86 | 4.87 | 1.12 | 3.75 | 6.37 | 1.69 | 1.31 | 10.3 | 3.93 | 5.99 | 11.61 | 5.06 | 3.75 | 1.12 | 3.75 | 534 |
| SOX-8Bm | 8.47 | 2.39 | 3.78 | 5.18 | 1 | 10.96 | 4.08 | 1.59 | 4.08 | 8.67 | 0.6 | 1.79 | 12.25 | 5.68 | 9.26 | 7.87 | 5.48 | 3.39 | 1.49 | 1.99 | 1004 |
| SOX-9BiBt | 8.02 | 0.38 | 4.2 | 5.92 | 2.1 | 6.49 | 4.01 | 2.1 | 4.77 | 4.77 | 2.29 | 2.67 | 16.98 | 8.21 | 4.01 | 9.35 | 4.96 | 4.2 | 0.95 | 3.63 | 524 |
| SOX-9Bt | 8.02 | 0.38 | 4.2 | 5.92 | 2.1 | 6.49 | 4.01 | 2.1 | 4.77 | 4.77 | 2.29 | 2.67 | 16.98 | 8.21 | 4.01 | 9.35 | 4.96 | 4.2 | 0.95 | 3.63 | 524 |
| SOX-10BiBt | 7.89 | 0.64 | 5.12 | 6.18 | 1.71 | 11.94 | 4.69 | 2.13 | 5.12 | 6.18 | 1.92 | 1.92 | 11.3 | 5.12 | 4.26 | 10.66 | 3.84 | 4.48 | 1.07 | 3.84 | 469 |
| SOX-10Bm | 8.32 | 0.44 | 5.03 | 5.03 | 1.75 | 11.6 | 4.81 | 2.41 | 5.47 | 5.91 | 1.97 | 1.97 | 11.38 | 5.69 | 4.6 | 9.85 | 4.6 | 4.16 | 1.09 | 3.94 | 457 |
| SOX-7BiBt | 8.53 | 1.55 | 4.91 | 5.94 | 1.81 | 9.3 | 3.36 | 0.78 | 3.62 | 10.59 | 2.33 | 2.58 | 12.4 | 4.13 | 5.94 | 10.34 | 3.36 | 3.36 | 0.78 | 4.39 | 387 |
| SOX-7Bm | 8.84 | 1.61 | 4.82 | 5.22 | 1.2 | 8.43 | 3.21 | 0.4 | 4.42 | 10.84 | 2.41 | 2.41 | 10.44 | 5.22 | 5.62 | 10.84 | 4.02 | 4.02 | 0.4 | 5.62 | 249 |
| SOX-17BiBt | 13.17 | 1.46 | 4.63 | 6.1 | 2.44 | 11.95 | 3.17 | 0.24 | 3.41 | 9.02 | 3.17 | 1.22 | 16.34 | 3.41 | 5.37 | 5.37 | 1.22 | 3.9 | 0.73 | 3.66 | 410 |
| SOX-17Bm | 11.57 | 1.85 | 3.7 | 7.41 | 2.78 | 10.65 | 2.78 | 0.93 | 3.7 | 8.33 | 1.85 | 3.24 | 12.04 | 1.85 | 9.26 | 7.41 | 2.31 | 4.63 | 1.39 | 2.31 | 216 |
| SOX-18BiBt | 14.65 | 1.8 | 4.11 | 6.68 | 2.57 | 8.23 | 1.03 | 0.77 | 2.83 | 9.51 | 1.29 | 1.54 | 19.28 | 2.06 | 8.74 | 5.91 | 2.06 | 2.83 | 1.03 | 3.08 | 389 |
| SOX-15BiBt | 7.3 | 0.86 | 2.15 | 4.29 | 1.72 | 11.16 | 2.58 | 0.43 | 4.72 | 7.73 | 2.15 | 3.43 | 14.59 | 5.15 | 6.87 | 12.02 | 3 | 3.43 | 1.72 | 4.72 | 233 |
| SOX-15Bm | 7.3 | 0.86 | 2.15 | 4.29 | 1.72 | 11.16 | 2.58 | 0.43 | 4.72 | 7.73 | 2.15 | 3.43 | 14.59 | 5.15 | 6.87 | 11.59 | 3.43 | 3.43 | 1.72 | 4.72 | 233 |
| SOX-30BiBt | 9.53 | 1.36 | 1.75 | 7.59 | 1.95 | 5.45 | 1.75 | 4.28 | 5.64 | 9.92 | 1.36 | 2.72 | 14.4 | 5.45 | 7.39 | 6.61 | 5.06 | 5.84 | 0.58 | 1.36 | 514 |
| SOX-30Bm | 8.13 | 1.26 | 2.66 | 7.15 | 2.81 | 5.19 | 3.09 | 3.93 | 4.35 | 8.42 | 0.98 | 3.09 | 13.88 | 4.91 | 6.17 | 9.82 | 5.47 | 5.33 | 0.42 | 2.95 | 713 |
| Avg | 9.86 | 1.07 | 4.21 | 6.26 | 1.78 | 8.72 | 3.08 | 2.39 | 5.32 | 7.79 | 2.73 | 2.74 | 10.97 | 5.76 | 5.82 | 9.6 | 4.1 | 3.95 | 0.89 | 2.97 | 473.2 |

**Figure 7:**
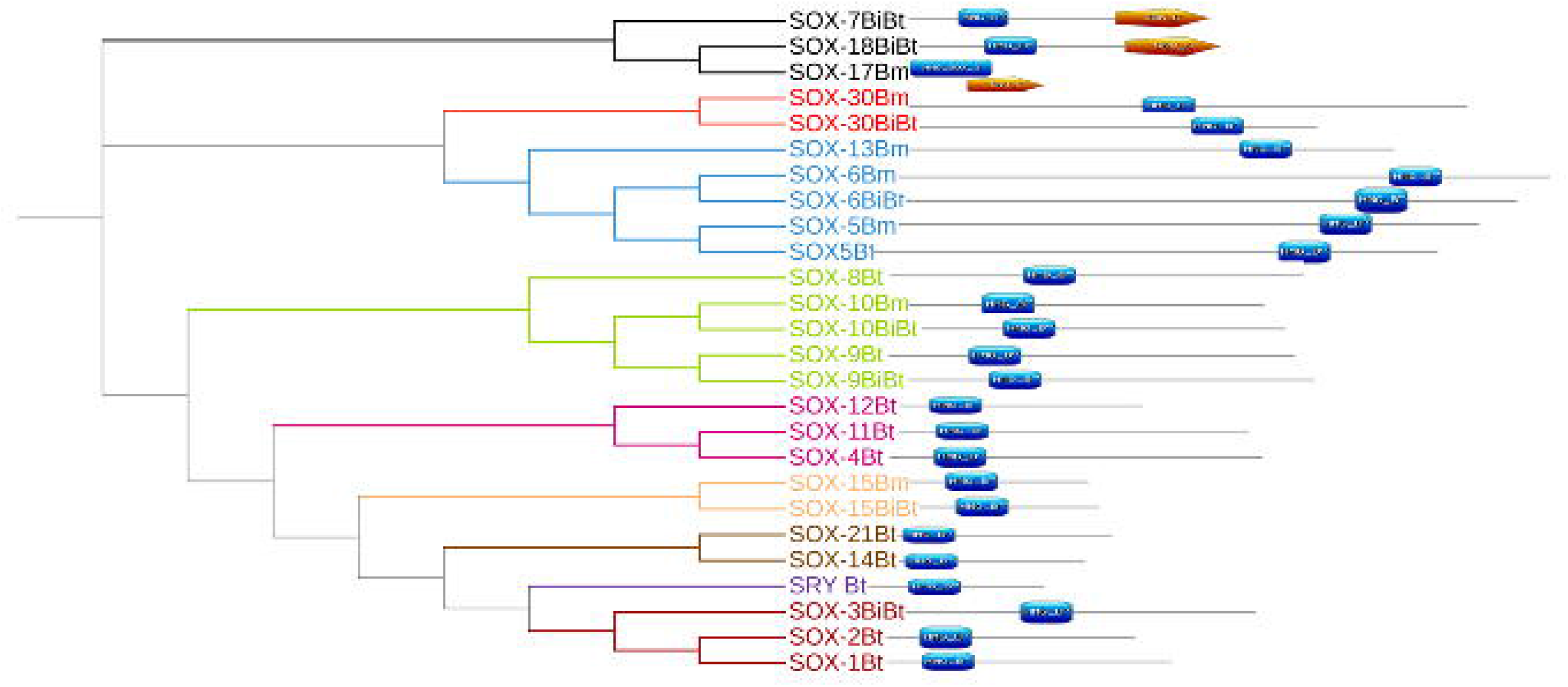
Comparison of intra-domain features of Sox proteins. This comparison shows the HMG box and SoxC domain which provides additional information about the structure and functions of Soc proteins. Members of SoxF subfamily have an additional C-terminal domain

### Biological, molecular, and cellular function of Bos Sox genes

For the Gene Ontology analysis, three criteria were utilized: the biological processes the genes were involved in, the cellular components they are part of, and their molecular function. We identified 49 biological processes with a false discovery rate >100, 6 cellular components, and 19 molecular functions (Table 4, Figure 8). DAVID analysis revealed 26 biological processes, 2 cellular components, and 7 molecular functions (Supplementary Table 1). Nineteen significant biological processes namely stem cell fate specification, positive regulation of mesenchymal cell, renal vesicle induction, retinal rod cell differentiation, metanephric nephron tubule formation, positive regulation of mesenchymal stem cell differentiation, endocardium formation, astrocytes fate commitment, endocardial cell differentiation, lacrimal gland development, ureter morphogenesis, Sertoli cell development, negative regulation of photoreceptor cell differentiation, regulation of stem cell proliferation, limb bud formation, enteric nervous system development, positive regulation of branching involved in ureteric bud morphogenesis, oligodendrocyte differentiation, negative regulation of myoblast differentiation and positive regulation of chondrocyte differentiation, were concomitantly identified from both databases.

**Table 4.** Gene Ontology biological processes, molecular functions, and cellular components of Bos Sox genes

| GO biological process complete | Gene | Pvalue | FDR |
| --- | --- | --- | --- |
| metanephric nephron tubule formation (GO:0072289) | 2 | 3.22E-06 | 5.59E-04 |
| renal vesicle induction (GO:0072034) | 2 | 3.22E-06 | 5.52E-04 |
| positive regulation of mesenchymal stem cell differentiation (GO:2000741) | 2 | 5.37E-06 | 8.39E-04 |
| endocardium formation (GO:0060214) | 2 | 5.37E-06 | 8.30E-04 |
| metanephric tubule formation (GO:0072174) | 2 | 5.37E-06 | 8.22E-04 |
| stem cell fate specification (GO:0048866) | 2 | 5.37E-06 | 8.13E-04 |
| astrocyte fate commitment (GO:0060018) | 2 | 8.05E-06 | 1.16E-03 |
| endocardial cell differentiation (GO:0060956) | 2 | 8.05E-06 | 1.15E-03 |
| cardiac endothelial cell differentiation (GO:0003348) | 2 | 8.05E-06 | 1.14E-03 |
| endocardium morphogenesis (GO:0003160) | 2 | 8.05E-06 | 1.12E-03 |
| metanephric nephron tubule morphogenesis (GO:0072282) | 2 | 1.13E-05 | 1.49E-03 |
| lacrimal gland development (GO:0032808) | 2 | 1.13E-05 | 1.47E-03 |
| common bile duct development (GO:0061009) | 2 | 1.13E-05 | 1.46E-03 |
| metanephric tubule morphogenesis (GO:0072173) | 2 | 1.50E-05 | 1.86E-03 |
| stem cell fate commitment (GO:0048865) | 2 | 1.50E-05 | 1.85E-03 |
| positive regulation of kidney development (GO:0090184) | 2 | 1.50E-05 | 1.83E-03 |
| negative regulation of photoreceptor cell differentiation (GO:0046533) | 2 | 1.50E-05 | 1.81E-03 |
| regulation of photoreceptor cell differentiation (GO:0046532) | 2 | 1.50E-05 | 1.80E-03 |
| regulation of mesenchymal stem cell differentiation (GO:2000739) | 2 | 1.93E-05 | 2.27E-03 |
| ureter morphogenesis (GO:0072197) | 2 | 1.93E-05 | 2.26E-03 |
| Sertoli cell development (GO:0060009) | 2 | 2.41E-05 | 2.69E-03 |
| endocardium development (GO:0003157) | 2 | 2.41E-05 | 2.67E-03 |
| limb bud formation (GO:0060174) | 2 | 2.94E-05 | 3.11E-03 |
| glial cell fate commitment (GO:0021781) | 2 | 3.53E-05 | 3.63E-03 |
| ureter development (GO:0072189) | 3 | 2.92E-07 | 6.57E-05 |
| metanephric nephron tubule development (GO:0072234) | 2 | 4.17E-05 | 4.20E-03 |
| positive regulation of chondrocyte differentiation (GO:0032332) | 3 | 4.08E-07 | 8.75E-05 |
| metanephric tubule development (GO:0072170) | 2 | 4.87E-05 | 4.83E-03 |
| sex determination (GO:0007530) | 3 | 4.75E-07 | 1.01E-04 |
| metanephric nephron epithelium development (GO:0072243) | 2 | 5.61E-05 | 5.53E-03 |
| retinal rod cell differentiation (GO:0060221) | 2 | 5.61E-05 | 5.49E-03 |
| positive regulation of branching involved in ureteric bud morphogenesis (GO:0090190) | 2 | 5.61E-05 | 5.45E-03 |
| enteric nervous system development (GO:0048484) | 2 | 5.61E-05 | 5.42E-03 |
| male sex determination (GO:0030238) | 2 | 5.61E-05 | 5.38E-03 |
| Sertoli cell differentiation (GO:0060008) | 2 | 6.41E-05 | 6.11E-03 |
| regulation of branching involved in ureteric bud morphogenesis (GO:0090189) | 2 | 7.26E-05 | 6.83E-03 |
| positive regulation of stem cell differentiation (GO:2000738) | 2 | 8.17E-05 | 7.58E-03 |
| metanephric nephron morphogenesis (GO:0072273) | 2 | 9.12E-05 | 8.41E-03 |
| metanephric epithelium development (GO:0072207) | 2 | 9.12E-05 | 8.36E-03 |
| nephron tubule formation (GO:0072079) | 2 | 1.01E-04 | 9.23E-03 |
| positive regulation of cartilage development (GO:0061036) | 3 | 1.45E-06 | 2.85E-04 |
| negative regulation of myoblast differentiation (GO:0045662) | 2 | 1.35E-04 | 1.20E-02 |
| regulation of stem cell proliferation (GO:0072091) | 3 | 2.53E-06 | 4.45E-04 |
| regulation of kidney development (GO:0090183) | 2 | 1.47E-04 | 1.30E-02 |
| metanephros morphogenesis (GO:0003338) | 2 | 1.60E-04 | 1.38E-02 |
| oligodendrocyte differentiation (GO:0048709) | 4 | 7.00E-08 | 1.74E-05 |
| metanephric nephron development (GO:0072210) | 2 | 2.01E-04 | 1.64E-02 |
| positive regulation of morphogenesis of an epithelium (GO:1905332) | 2 | 2.01E-04 | 1.63E-02 |
| developmental induction (GO:0031128) | 2 | 2.01E-04 | 1.62E-02 |
| **GO molecular function complete** |  |  |  |
| beta-catenin binding (GO:0008013) | 3 | 2.65E-05 | 6.61E-03 |
| DNA-binding transcription activator activity, RNA polymerase II-specific (GO:0001228) | 8 | 3.12E-10 | 8.76E-08 |
| DNA-binding transcription activator activity (GO:0001216) | 8 | 3.36E-10 | 8.89E-08 |
| RNA polymerase II cis-regulatory region sequence-specific DNA binding (GO:0000978) | 18 | 9.28E-25 | 4.17E-21 |
| cis-regulatory region sequence-specific DNA binding (GO:0000987) | 18 | 1.32E-24 | 2.98E-21 |
| DNA-binding transcription factor activity, RNA polymerase II-specific (GO:0000981) | 18 | 6.41E-24 | 9.61E-21 |
| RNA polymerase II transcription regulatory region sequence-specific DNA binding (GO:0000977) | 18 | 1.06E-23 | 1.20E-20 |
| DNA-binding transcription factor activity (GO:0003700) | 18 | 1.72E-23 | 1.54E-20 |
| transcription cis-regulatory region binding (GO:0000976) | 18 | 3.77E-23 | 2.83E-20 |
| transcription regulatory region nucleic acid binding (GO:0001067) | 18 | 3.82E-23 | 2.46E-20 |
| sequence-specific double-stranded DNA binding (GO:1990837) | 18 | 7.82E-23 | 4.40E-20 |
| double-stranded DNA binding (GO:0003690) | 18 | 2.44E-22 | 1.22E-19 |
| sequence-specific DNA binding (GO:0043565) | 18 | 2.78E-22 | 1.25E-19 |
| transcription regulator activity (GO:0140110) | 18 | 3.37E-21 | 1.38E-18 |
| DNA binding (GO:0003677) | 18 | 5.82E-19 | 2.18E-16 |
| nucleic acid binding (GO:0003676) | 18 | 2.25E-15 | 7.79E-13 |
| heterocyclic compound binding (GO:1901363) | 18 | 1.08E-11 | 3.48E-09 |
| organic cyclic compound binding (GO:0097159) | 18 | 1.38E-11 | 4.15E-09 |
| binding (GO:0005488) | 18 | 3.38E-05 | 8.00E-03 |
| **GO cellular component complete** |  |  |  |
| transcription regulator complex (GO:0005667) | 6 | 7.39E-07 | 6.94E-04 |
| nucleus (GO:0005634) | 18 | 2.33E-10 | 4.38E-07 |
| intracellular membrane-bounded organelle (GO:0043231) | 18 | 1.57E-06 | 9.86E-04 |
| membrane-bounded organelle (GO:0043227) | 18 | 3.34E-06 | 1.57E-03 |
| intracellular organelle (GO:0043229) | 18 | 1.51E-05 | 5.69E-03 |
| organelle (GO:0043226) | 18 | 3.48E-05 | 1.09E-02 |

**Figure 8:**
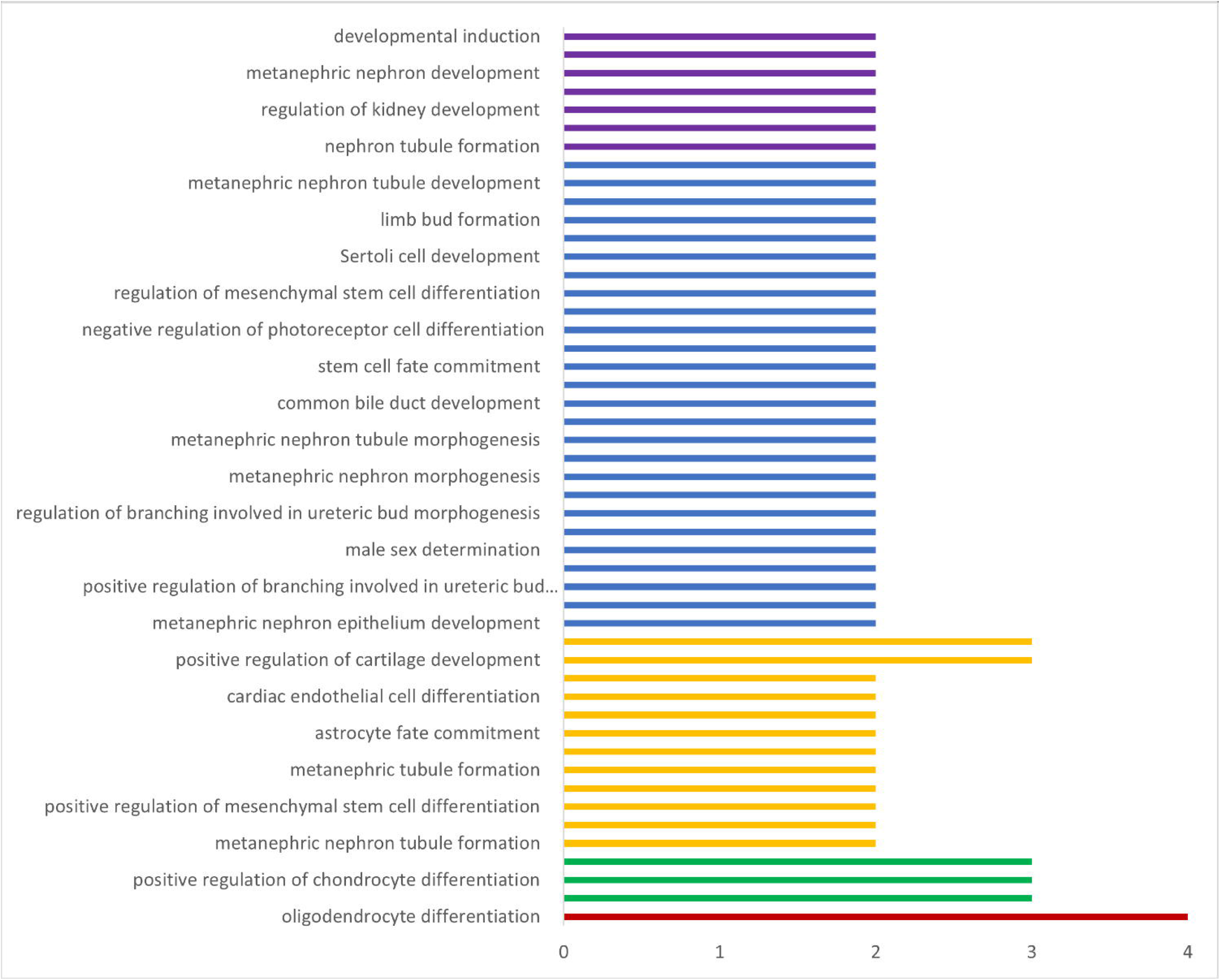
The biological process and the corresponding number of target genes involved. Red bars indicate the biological process/pathways predicted with the highest *p*-value, purple bars indicate the pathways predicted with the lowest *p*-value

### Protein-protein interaction cluster with Sox genes

In order to analyze the protein-protein interaction, co-expression, genetic interaction, and physical association, we used STRING to build the protein network. Using k-means clustering we generated 3 clusters composed of closely connected interactions. The results revealed a major PPI network cluster with all the studied genes except Sox12, 15, and 30, which exhibited no interaction (Figure 9a). Sox5 had the most interaction with 10 nodes while Sox11, 13, and 14 had the least interaction. However, Sox1, 2, and 3 were not found in the cluster.

**Figure 9:**
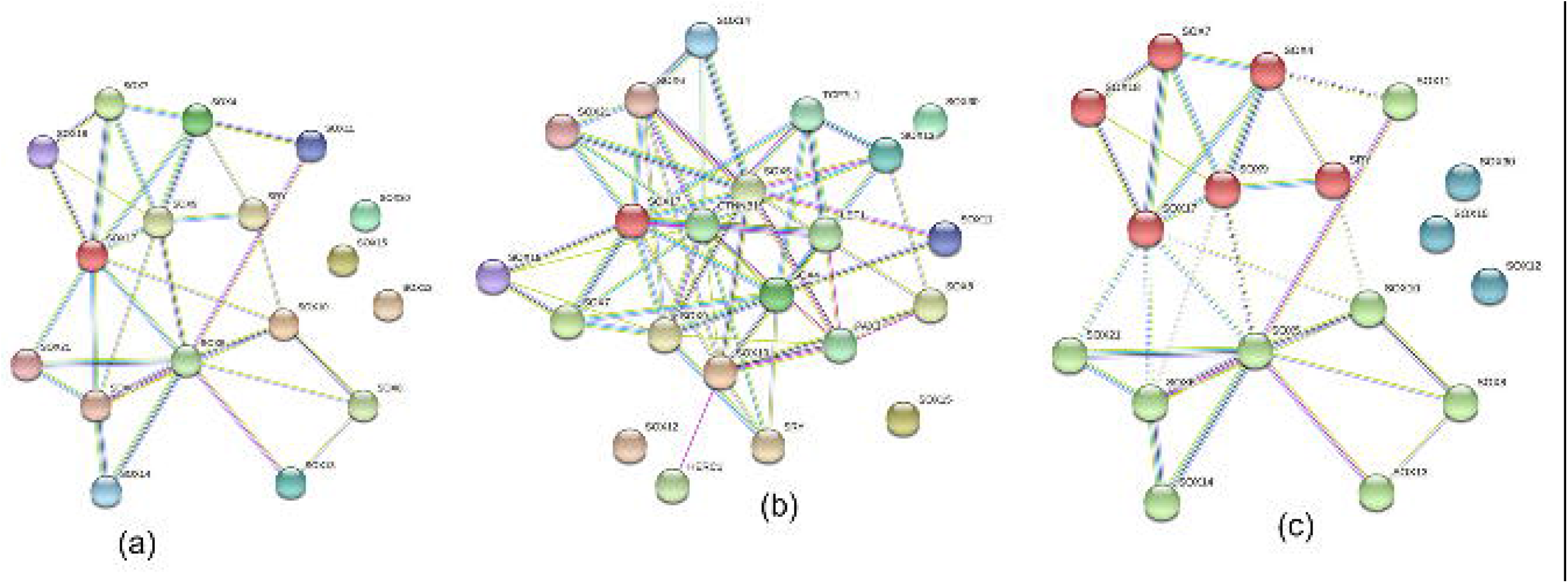
Sox protein interaction. Protein protein interaction (a) Sox proteins only (b) predicted interactions with other proteins (c) k means clustering. Network created using STRING (string-db.org)

The protein-protein interaction analysis indicated the genes’ possible involvement in pathways such as metabolic, developmental, and regulatory. Cellular signaling pathways such as regulation of Wnt signaling pathway, negative regulation of canonical Wnt signaling pathway, regulation of canonical Wnt signaling pathway, and signal transduction involved in cell cycle checkpoint were associated with these protein network. STRING extended analysis reveal the interaction of specific Sox genes with CTNNB1, HERC1, LEF1, PAX3, and TCF7L1 (Figure 9b).

## Discussion

Sox family of transcription factors exhibit diverse tissue-specific expression patterns during early embryonic development and plays important roles in cell fate (Bowles et al., 2000). To provide a deeper understanding of the role of *Sox* protein in Bovidae evolution, we utilized sequences available in public databases to perform computational analysis, yielding evolutionary relationships among Sox gene family in Bos. Our analysis identified nine distinct clades, which were assigned to the known Sox groups A -H (Jiang et al., 2019; Hu et. al., 2021). The genes within the same subgroup shared greater sequence similarity outside the HMG domain than those that are more related, a finding consistent with previous report (Heenan et al., 2016). We observed high percentages of proline, glycine, and alanine in all the Sox proteins. Studies have shown that proline readily adopts a *cis* and *trans* configuration in response to subtle influences, induces sharp turns in the local geometry and is important in protein-protein interaction, cell adhesion, and mediates signal transduction (Morgan and Rubenstein, 2013).

SoxG and SoxD subfamily had the lowest and highest amino acid residues and molecular weights respectively. All the Sox proteins had negative GRAVY values, indicating the hydrophilic nature of the proteins, as previously described (Jiang et al., 2019; Zhang et al., 2018), enhancing their binding capacity (Roy et al., 2011: Kadel et al., 2019) SoxA (*SRY*) protein, a transcriptional regulator that controls the genetic switch in male development, is the only member of Group A and formed a monophyletic group in the phylogenetic analysis. Its expression inhibition cause defects in testes development (Larney et. al., 2014, Hu et al., 2021), with cytogenetic and molecular studies revealing it possibly arose from a mutation of Sox 3 and fusion of the gene with regulatory sequences from another gene already on the X chromosome (Herpin and Schart, 2015). STRING pathway analysis revealed its association with Sox 9, with GO enrichment confirming that SRY and Sox9 are enriched in male sex determination, which is consistent with published reports (Sun et al., 2014; Jiang et.al., 2019). *SRY* binds to TESCO (testis-specific enhancer core) sequence of Sox9 and initiates the differentiation of somatic precursors into Sertoli cells that coordinate the gonadal development toward the testis (Wilhelm et al., 2007). Despite substantial variation in expression profile structure and amino acid sequences within mammals, the function of *SRY* to activate Sox9 during development appears to be conserved.

The Sox B_1_ group comprises Sox1, 2, and 3, groups of transcription factors that have been reported to affect Wnt β-catenin signaling pathway (Stevanovic et. al., 2021). Sox1 inhibits β-catenin/TCF transcription activity binding to β-catenin via their C-terminal region while Sox2 overexpression down-regulates the Wnt β-catenin signaling pathway suggesting a negative feedback loop between them. These proteins also promote the self-renewal of neural progenitor cells. PANTHER analysis shows that Sox3 interacts with *SRY* and Sox9 in sex determination. There was no interaction reported for Sox1 and 2, possibly attributed to functional redundancy in this subfamily (Brunelli et al., 2003; Weiss et al., 2003)

Proteins encoded by members of SoxB_2_ subfamily: Sox14 and Sox21 are very similar to each other but different from subfamily B_1_ in areas outside the HMG domain. They have been implicated in the regulation of mesenchymal stem cell differentiation. Sox21 has been reported to promote neurogenesis in mice while Sox14 mediates terminal differentiation of the neural progenitor cells (Makrides et al., 2018). STRING pathway analysis identified Sox14 as one of the genes involved in the regulation of neurogenesis while Sox21 was enriched in stem cell differentiation.

SoxC subfamily (Sox4, 11, and 12) has three members in most vertebrates (Dy et al., 2008). Sox4 and 11 share similar function and structural properties. Reports on Sox12 functional properties are scarce. Sox4 and 11 were shown to be involved in lymphocyte differentiation, osteoblast development, neural and glial cell development, control progenitor development, and capacity to crosstalk with other cells to grow and mature skeletal structures (Dy et al., 2008, Lefebvre, 2019). Phylogenetic analysis reveals that the three proteins are closely related, as they all appeared in one clade. PROSITE revealed a SoxC-rich domain in all SoxF genes suggesting that the complex interacts with this group. The HMG box domain in this group is virtually identical, therefore the difference in DNA binding efficiency may be due to sequence differences outside the HMG box (Dy et al., 2008). STRING pathway analysis identified Sox4 and Sox11 as enriched in cardiac ventricle formation and limb bud formation, enteric nervous system development, and glial cell proliferation.

SoxD subfamily (Sox5, 6 and 13) have higher amounts of amino acid residues than other Sox proteins. Our result showed that this subgroup is significantly enriched in oligodendrocyte differentiation, cell fate commitment, and glial cell differentiation. Members of this group formed a paraphyletic group with Sox30 suggesting a recent divergence. These genes interact with Sox9 of the SoxE subfamily, and expression of the three Sox genes culminates in growth plate proliferation in chondrocytes Furthermore, co-inactivation of Sox5 and 6 genes was reported to result in stunted or underdeveloped growth plates and articular cartilage, while Sox9 is required to turn on and maintain chondrocyte-specific genes (Han and Lefebvre, 2008. Lui and Lefebvre, 2015).

SoxE proteins, Sox8, 9, and 10, overlap in expression, play critical roles in various biological processes and are renowned for their role in transactivation. Haseeb and Lefebvre (2019) reported that SoxE and F proteins possess the unique motif the EFDQYL/ELDQYL required for transactivation. This motif is functionally crucial and highly conserved in various vertebrates and invertebrates species and is categorized based on amino acid composition (Frietze and Farnham, 2011). The residues within the motif revealed the presence of acidic and hydrophilic amino acids (E, Q) alternating with hydrophobic residues (F, Y, L) which fold into pocket-like structures involved in recognizing and binding to functional partners and coactivators (Haseeb and Lefebvre, 2019). The identified motif was found in all SoxE proteins in Bos sp. Previous studies showed that the sequence is required for the interaction of Sox9 and β-catenin, (Akiyama et. al., 2004)

The SoxF subfamily promotes fate specification of endothelial cells (Kim et al., 2016). They are thought to share a recent common ancestor with SoxE and exhibit substantial overlap in expression function and regulation implying a redundant type of cooperation. We found this EFDQYL/ELDQYL motif in Sox7 and Sox 18 but not in Sox17. The only Sox member in the G subfamily is Sox 15. This Sox protein is found only in mammals and share a common ancestry with SoxB, forming a paraphyletic tree. (Kamachi and Kondoh, 2013; Ito, 2010; Williams et al., 2020). This common ancestry likely accounts for the ability of Sox15 to replace the function of Sox2 in the self-renewal of mouse stem cells (Maruyama et al., 2005; Niwa et al., 2016)

STRING analysis revealed complex interaction of the studied Sox proteins with CTNNB1, PAX3, HERC1, and TCF7L1. Catenin γ-1 gene (CTNNB1) provides instructions for making the beta-catenin protein which is primarily found at junctions connecting neighboring cells. Beta-catenin plays a major role in sticking cells (adhesion), cell signaling, and in cell communication (Bienz, 2005). This pathway is important because cellular interactions are crucial for Sox proteins to recognize target genes. Jiang et al, 2019 reported that CTNNB1 formed complex interactions with Sox 2,6,7 and 9 in several developmental processes in *Coturnix. japonica*. STRING analysis also revealed the complex interaction of CTNNB1, Sox 17, 30, LEF1, and TCF7L1 in beta-catenin binding. LEF1 (lymphoid enhancer-binding factor 1) shares significant homology with HMG protein and are involved in Wnt canonical pathway, cell differentiation, and follicle morphogenesis (Xiao et al., 2021). Pax3, Sox10, and c-Ret are components of a neural crest development pathway, and interruption of this pathway at various stages results in neural crest–related human genetic syndromes (Lang et al., 2000)

## Conclusion

Our study shows a detailed molecular evolution, relationship and functions of Sox genes in the Bos family using several computational tools. Also, this study documents the first comprehensive genetic variation of the Sox gene family in Bos sp. We reported a total of 20 Sox genes which were classified into nine subfamilies based on phylogenetic analysis. We also provided evidence of the divergent evolution some Sox genes among the Bos family. This study further revealed the functional motifs that drive these proteins and enhances their ability to shape the regulatory regions of several genes and promote cell fate and differentiation. Lastly, we found the motif EFDQYL/ELDQYL which is required for the unique transactivation ability of SoxE and SoxF proteins. Based on the findings of this study we provide suitable backgrounds for further study to harness the potential of these protein family in immunotherapeutic and regenerative medicine targeted in Bovidae.

## Acknowledgement

OBM was supported by Pitt-Momentum Fund of the University of Pittsburgh. Ongoing support by the Division of Biological and Health Sciences, Pitt-Bradford, College of Health Sciences and Technology, Rochester Institute of Technology is acknowledged. APC charges for this article were fully paid by the University Library System, University of Pittsburgh.

## Author Contributions

OBM and MOA conceptualized and designed the experiments; MOA, MSA and OBM carried out the experiments, analyzed the data and drafted the manuscript; MOA, MSA, OHO, JF, AG, SOP, IMO, BNT and OBM revised the manuscript, contributed to the discussion and scientific content. All authors read and approved the final version of the manuscript.

## Data Availability (see supplementary Tables 1)

**Supplementary Figure 1**. EFDQYL and ELDQYL conserved domain in SoxE and SoxF subfamilies

**Supplementary Table 1:** Genetic Distance of Bos sp Sox genes subfamilies

**Supplementary Table 2: DAVID Analysis Output**

